# Improved multi-type birth-death phylodynamic inference in BEAST 2

**DOI:** 10.1101/2020.01.06.895532

**Authors:** Jérémie Scire, Joëlle Barido-Sottani, Denise Kühnert, Timothy G. Vaughan, Tanja Stadler

## Abstract

The multi-type birth-death model with sampling is a phylodynamic model which enables quantification of past population dynamics in structured populations, based on phylogenetic trees. The BEAST 2 package *bdmm* implements an algorithm for numerically computing the probability density of a phylogenetic tree given the population dynamic parameters under this model. In the initial release of *bdmm*, analyses were limited to trees consisting of up to approximately 250 genetic samples for numerical reasons. We implemented important algorithmic changes to *bdmm* which dramatically increase the number of genetic samples that can be analyzed, and improve the numerical robustness and efficiency of the calculations. Being able to use bigger datasets leads to improved precision of parameter estimates. Furthermore, we report on several model extensions to *bdmm*, inspired by properties common to empirical datasets. We apply this improved algorithm to two partly overlapping datasets of Influenza A virus HA sequences sampled around the world, one with 500 samples, the other with only 175, for comparison. We report and compare the global migration patterns and seasonal dynamics inferred from each dataset.

**Availability:** The latest release with our updates, *bdmm* 0.3.5, is freely available as an open access package of BEAST 2. The source code can be accessed at *https://github.com/denisekuehnert/bdmm*.

## Introduction

Genetic sequencing data taken from a measurably evolving population contain fingerprints of past population dynamics [Felsenstein, 1992]. In particular, the phylogeny spanning the sampled genetic data contains information about the mixing pattern of different populations and thus contains information beyond what is encoded in classic occurrence data, see e.g. Hey and Machado [2003], Stadler and Bonhoeffer [2013b]. Phylodynamic methods [Grenfell et al., 2004, Kühnert et al., 2011] aim at quantifying past population dynamic parameters, such as migration rates, from genetic sequencing data. Such tools have been widely used to study the spread of infectious diseases in structured populations, see e.g. Dudas et al. [2017], Faria et al. [2018] as examples for analyses of recent epidemic outbreaks. Both the host population and the pathogen population may be structured, e.g. the host population may be geographically structured, and the pathogen population may consist of a drug-sensitive and a drug-resistant subpopulation. Understanding how these subpopulations interact with one another, whether they are separated by geographic distance, lifestyles of the hosts, or other barriers, is a key determinant in understanding how an epidemic spreads. In macroevolution, different species may also be structured into different “subpopulations”, e.g. due to their geographic distribution or to trait variations, see e.g. Hodges [1997]. Phylodynamic tools aim at quantifying the rates at which species migrate or traits are gained or lost, and the rates of speciation and extinction within the “subpopulations”, see e.g. Goldberg et al. [2010], Mayrose et al. [2011], Goldberg et al. [2011]. The phylodynamic analysis of structured populations can be performed using two classes of models, namely coalescent-based and birth-death-based approaches. Both have their unique advantages and disadvantages [Volz and Frost, 2014, Boskova et al., 2014]. Here, we report on improvements to a multi-type birth-death-based approach.

A multi-type birth-death model is a linear birth-death model accounting for structured populations. Under this model, the probability density of a phylogenetic tree can be calculated by numerically integrating a system of differential equations. The use of this model within a phylodynamic setting and the associated computational approach were initially proposed for analyzing species phylogenies [Maddison et al., 2007] and later for analyzing pathogen phylogenies [Stadler and Bonhoeffer, 2013a, Volz and Frost, 2014]. The package *bdmm* within the Bayesian phylodynamic inference framework BEAST2 [Bouckaert et al., 2014] generalizes the assumptions of these two initial approaches [Kühnert et al., 2016]. It further allows for co-inferring phylogenetic trees together with the model parameters and thus takes phylogenetic uncertainty explicitly into account. Datasets containing more than 250 genetic sequences could not be analysed using the original *bdmm* package [Kühnert et al., 2016] due to numerical instability. This limitation was a strong impediment to obtaining reliable results, particularly for analysis of structured populations, as quantifying parameters which characterize each subpopulation requires a significant amount of samples from each of them. The instability was due to numerical underflow in the probability density calculations, meaning that probability values extremely close to zero could not be accurately calculated and stored. We have solved the numerical instability issue of *bdmm*, thereby lifting the hard limit on the number of samples that can be analysed. In addition, the practical usefulness of the *bdmm* package was previously restricted by the amount of computation time required to carry out analyses. We report here on significant improvements in computation efficiency. As a result, *bdmm* can now handle datasets containing several hundred genetic samples. Finally, we made the multi-type birth-death model more general in several ways: homochronous sampling events can now occur at multiple times (not only the present), individuals are no longer necessarily removed upon sampling, and the migration rate specification has been made more flexible by allowing for piecewise-constant changes through time.

Overall, these model generalizations and implementation improvements enable more reliable and ambitious empirical data analyses. Below, we use the new release of *bdmm* to quantify Influenza A virus spread around the globe as an example application, and compare the results obtained with those from the reduced dataset analysed in [Kühnert et al., 2016].

## Methods

### Description of the extended multi-type birth-death model

First, we formally define the multi-type birth-death model on d types [Kühnert et al., 2016] including the generalizations introduced in this work. The process starts at time 0 with one individual, this is also called the origin of the process and the origin of the resulting tree. This individual is of type *i* ∈ {1…*d*}, with probability *h*_*i*_ (where 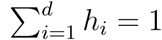). The process ends after *T* time units (at present). The time interval (0, *T*) is partitioned into *n* intervals through 0 < *t*_1_ < … < *t*_*n*−1_ < *T*, and we define *t*_0_ := 0 and *t*_*n*_ := *T*. Each individual at time *t, t*_*k*−1_ ≤ *t* < *t*_*k*_, *k* ∈ {1… *n*} of type *i* ∈ {1… *d*}, gives rise to an additional individual of type *j* ∈ {1… *d*}, with birth rate *λ*_*ij,k*_, migrates to type *j* with rate *m*_*ij,k*_ (with *m*_*ii,k*_ = 0), dies with rate *μ*_*i,k*_, and is sampled with rate *ψ*_*i,k*_. At time *t*_*k*_, each individual of type *i* is sampled with probability *ρ*_*i,k*_. Upon sampling (either with rate *ψ*_*i,k*_ or probability *ρ*_*i,k*_), the individual is removed from the infectious pool with probability *r*_*i,k*_. We summarize all birth-rates *λ*_*ij,k*_ in ***λ***, migration rates *m*_*ij,k*_ in ***m***, death rates *μ*_*i,k*_ in ***μ***, sampling rates *ψ*_*i,k*_ in ***ψ***, sampling probabilities *ρ*_*i,k*_ in ***ρ*** and removal probabilities *r*_*i,k*_ in ***r***, *i, j* ∈ {1,…, *d*}, *k* ∈ {1,…, *n*}. The model described in Kühnert et al. [2016] is a special case of the above, assuming that migration rates are constant through time (i.e. do not depend on *k*), removal probabilities are constant through time and across types (i.e. do not depend on *k* and *i*), and that *ρ*_*i,k*_ = 0 for *k* < *n* and *i* ∈ {1… *d*}. This process gives rise to complete trees on sampled and non-sampled individuals with types being assigned to all branches at all times (Figure 1, left). Following each branching event, one offspring is assigned to be the “left” offspring, and one the “right” offspring, each assignment has probability 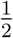. In the figure, we draw the branch with assignment “left” on the left and the branch with assignment “right” on the right. Such trees are called oriented trees, and considering oriented trees will facilitate calculations of probability densities of trees. Pruning all lineages without sampled descendants leads to the *sampled phylogeny* (Figure 1, middle and right). The orientation of a branch in the sampled phylogeny is the orientation of the corresponding branch descending the common branching event in the complete tree. When the sampled phylogeny is annotated with the types along each branch, we refer to it as a *branch-typed tree* (Figure 1, middle). On the other hand, if we discard these annotations but keep the types of the sampled individuals, we call the resulting object a *sample-typed (or tip-typed) tree* (Figure 1, right). Below, we state the probability density of the sampled tree (i.e. the sample-typed or branch-typed tree) given the multi-type birth–death parameters ***λ, m, μ, ψ, ρ, r***, *T*. This probability density is obtained by integrating probability densities g from the leaf nodes (or “tips”), backwards along all edges (or “branches”), to the origin of the tree. Our notation here is based on previous work [Kühnert et al., 2016, Stadler et al., 2013], and the probabilities *p*_*i,k*_(*t*) and 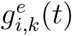 relate to *E* and *D* in Maddison et al. [2007], Stadler and Bonhoeffer [2013a], respectively.

**Figure 1:**
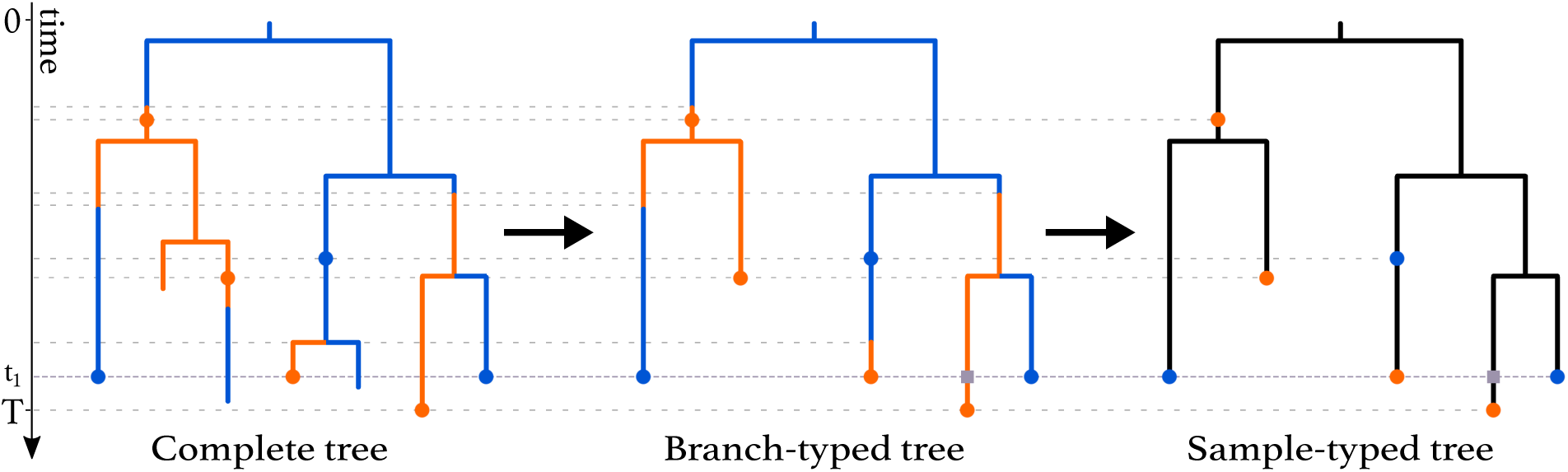
Complete tree (left) and sampled trees (middle and right) obtained from a multi-type birth-death process with two types. The orange and blue dots on the trees represent sampled individuals and are coloured according to the type these individuals belong to. A *ρ*-sampling event happens at time *t*_1_. The grey squares represent degree-2 nodes added to branches crossing this event. *ρ*-sampling also happens at present (time *T*). As seen in the complete tree, the first three individuals who were sampled were not removed from the population upon sampling, while the four individuals giving rise to the later samples were removed upon sampling.

Every branching event in the sampled tree gives rise to a node with degree 3 (i.e. 3 branches are attached). Every sampling event gives rise to a node of degree 2 (called “sampled ancestor”) or 1 (called “tip”, as defined above). A sampling event at time *t* = *t*_*k*_, *k* ∈ {1,…, *n*}, is referred to as a *ρ*-sampled node. All other nodes corresponding to samples are referred to as *ψ*-sampled nodes. Further, degree-2 nodes are put at time *t*_*k*_ on all lineages crossing time *t*_*k*_, *k* = 1,…, *n* − 1 as shown at time *t*_1_ in Figure 1. In a branch-typed tree, a node of degree 2 also occurs on a lineage at a time point when a type-change occurs. Such type changes may be the result of either migrations or birth events in which one of the descendant subtrees is unsampled (Figure 1, middle).

We highlight that in *bdmm*, we assume that the most recent sampling event happens at time *T*. This is equivalent to assuming that the sampling effort was terminated directly after the last sample was collected, and overcomes the necessity for users to specify the time between the last sample and the termination of the sampling effort at time *T*.

The derivation of the probability density of a sampled tree under the extended multi-type birth-death model is developped in Supplementary Information (SI) (section S1).

### Implementation improvements

The computation of probability densities of sampled trees under the multi-type birth-death model require numerically solving Ordinary Differential Equations (ODEs) along each tree branch. We significantly improved the robustness of the original *bdmm* implementation, which suffered from instabilities caused by underflow of these numerical calculations. Compared to the original implementation, we prevent underflow by implementing an extended precision floating point representation (EPFP) for storing intermediary calculation results. Additional to this improvement in stability, we improved the efficiency of the probability density calculations, by 1) using an adaptive-step-size integrator for numerical integration, 2) performing preliminary calculations and storing the results for use during the main calculation step and 3) distributing calculations among threads running in parallel. Details can be found in SI section S2.

## Results

### Evaluation of numerical improvements

We compared the robustness and efficiency of the improved *bdmm* package against its original version. We considered varying tree sizes, between 10 and 1000 samples. For each tree size, we simulated 50 branch-typed and 50 sample-typed trees under the multi-type birth-death model using randomly-drawn parameter values from the distributions shown in SI Table S1. For each simulated tree, we measured the time taken to perform the calculation of the probability density given the parameter values under which the tree was simulated, using the updated and the original *bdmm* implementation. We report the wall-clock time taken to perform this calculation 5000 times (Fig. 2). All computations are performed on a MacBook Pro with a dual-core 2.3 GHz Intel Core i5 processor. The new implementation of *bdmm* is on average 9 times faster than the original (Fig. 2A). The robustness of the updated implementation is demonstrated by reporting how often the implementations return −∞ for the probability density in log space. We call these calculations “failures”, the most likely cause of error being underflow. Our new implementation shows no calculation failure for trees with up to 1000 samples, while in the original implementation calculations often fail for trees with more than 250 samples (Fig. 2B).

**Figure 2:**
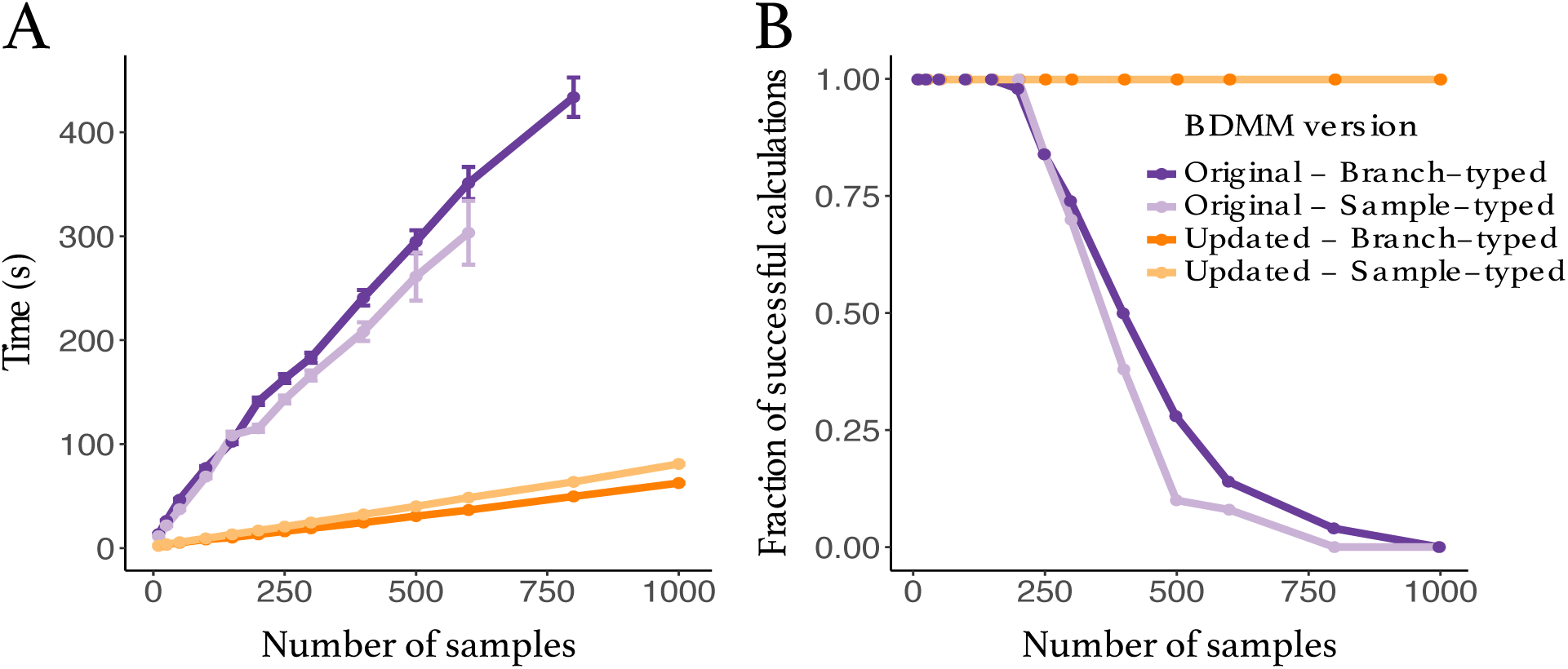
Comparison between the original and updated implementation of the multi-type birth-death model. **A:** Speed comparison. Only successful calculations were taken into account i.e. calculations where the log-probability density was different from −∞. **B:** Success in calculating probability density values plotted against tree size. The values presented in this panel correspond to the same set of calculations as the one in panel A.

### Influenza A virus (H3N2) analysis

As an example of a biological question which can be investigated with *bdmm*, we analysed 500 H3N2 influenza virus HA sequences sampled around the globe from 2000 to 2006 and aim to recover the seasonal dynamics of the global epidemics. The dataset is a subset of the data analysed by Vaughan et al. [2014], taken from three different regions around the globe: New York (North, *n* = 167), New Zealand (South, *n* = 215) and Hong Kong (Tropics, *n* = 118). As a comparison, we performed an identical analysis with the H3N2 influenza dataset of 175 sequences sampled between 2003 and 2006 used in [Kühnert et al., 2016]. This dataset of 175 sequences was also a subset of the data by Vaughan et al. [2014], and it also groups samples from 3 locations denoted as North (for northern hemisphere), South (for southern hemisphere) and Tropic (for tropical regions). Note that the latter dataset comes from more geographically-spread samples and thus we do not expect results from both analysis to be perfectly comparable. As we deal with pathogen sequence data, we adopt the epidemiological parametrization of the multi-type birth-death model as detailed in Kühnert et al. [2016]. The epidemiological parametrization substitutes birth, death and sampling rates with effective reproduction numbers within types, rate at which hosts become noninfectious and sampling proportions. To study the seasonal dynamics of the global epidemic, we allow the effective reproduction number *R*_*e*_ to vary through time. To do so, we subdivide time into six-month intervals (starting April 1st and October 1st) and we constrain effective reproduction number values corresponding to the same season across different years to be equal for each particular location. Further details on the data analysis configuration can be found in Supplementary section S3.

The analysis of the larger dataset (500 samples) allows for the reconstruction of the evolutionary tree encompassing a longer time period, and therefore gives a more long-term and detailed view of the evolution of the global epidemic (see Fig 3 for the Maximum-Clade Credibility trees). As can be expected for the tropical location, in both analyses, effective reproduction numbers for H3N2 influenza A are inferred to be close to one year-round (Fig 4A). Conversely, strong seasonal variations can be observed in Northern and Southern hemisphere locations. There, the effective reproduction number is close to one in winter, while it is much lower in summer. Inferences from the small and large datasets are mostly in agreement. A subtle difference appears: in the larger dataset, the effective reproduction number in winter seasons and in the tropical location are closer to one, with less variation across estimates. This seems to indicate that the variations between estimates observed with the smaller dataset including samples from 2003 to 2006 (for instance *R*_*e*_ in winter in the North compared to *R*_*e*_ in winter in the South) are due to stochastic fluctations which are averaged out when considering a longer period of transmission dynamics in the larger dataset covering the years 2000 to 2006.

**Figure 3:**
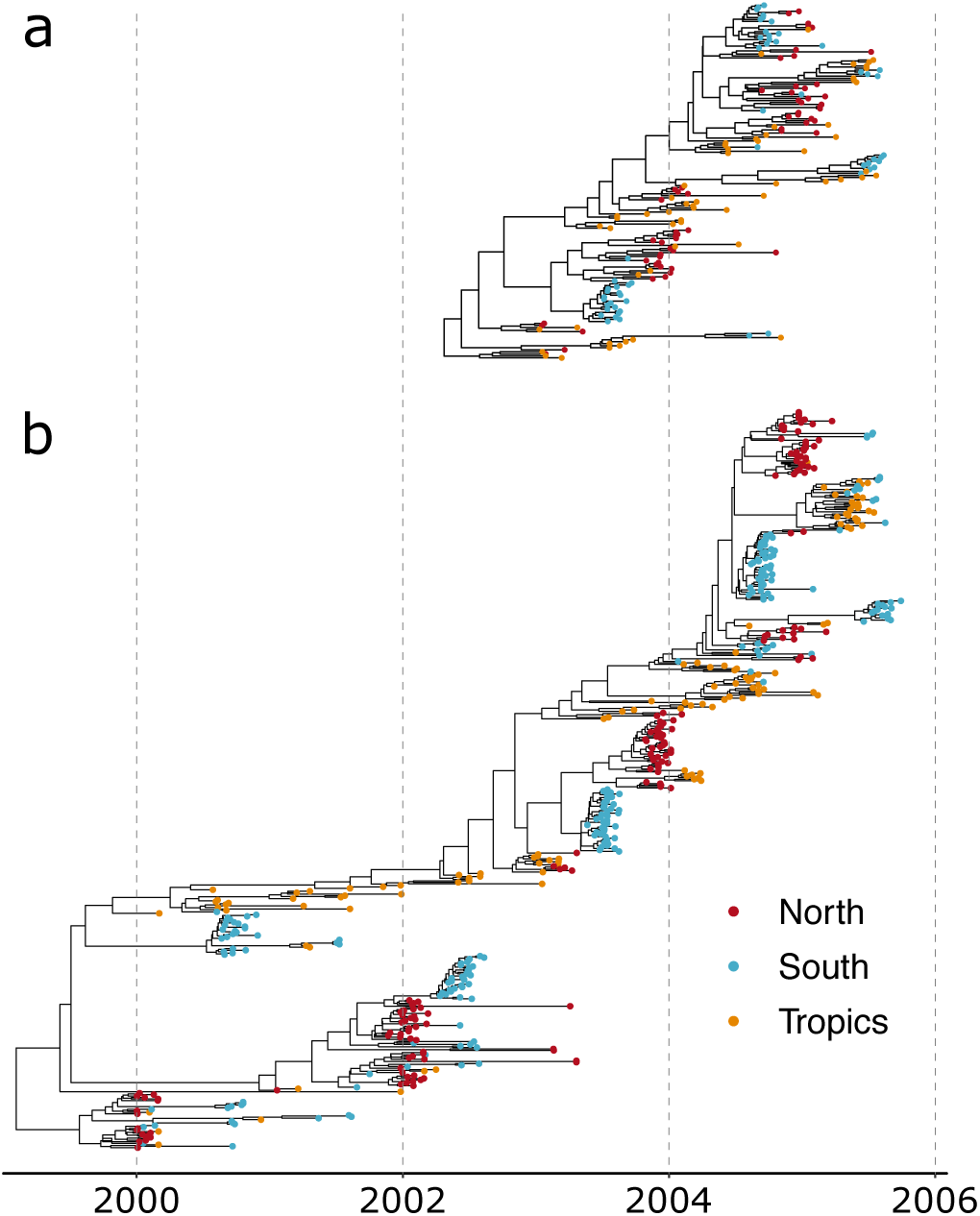
Maximum-Clade Credibility (MCC) trees for analyses with 175 samples (a) and 500 samples (b).

**Figure 4:**
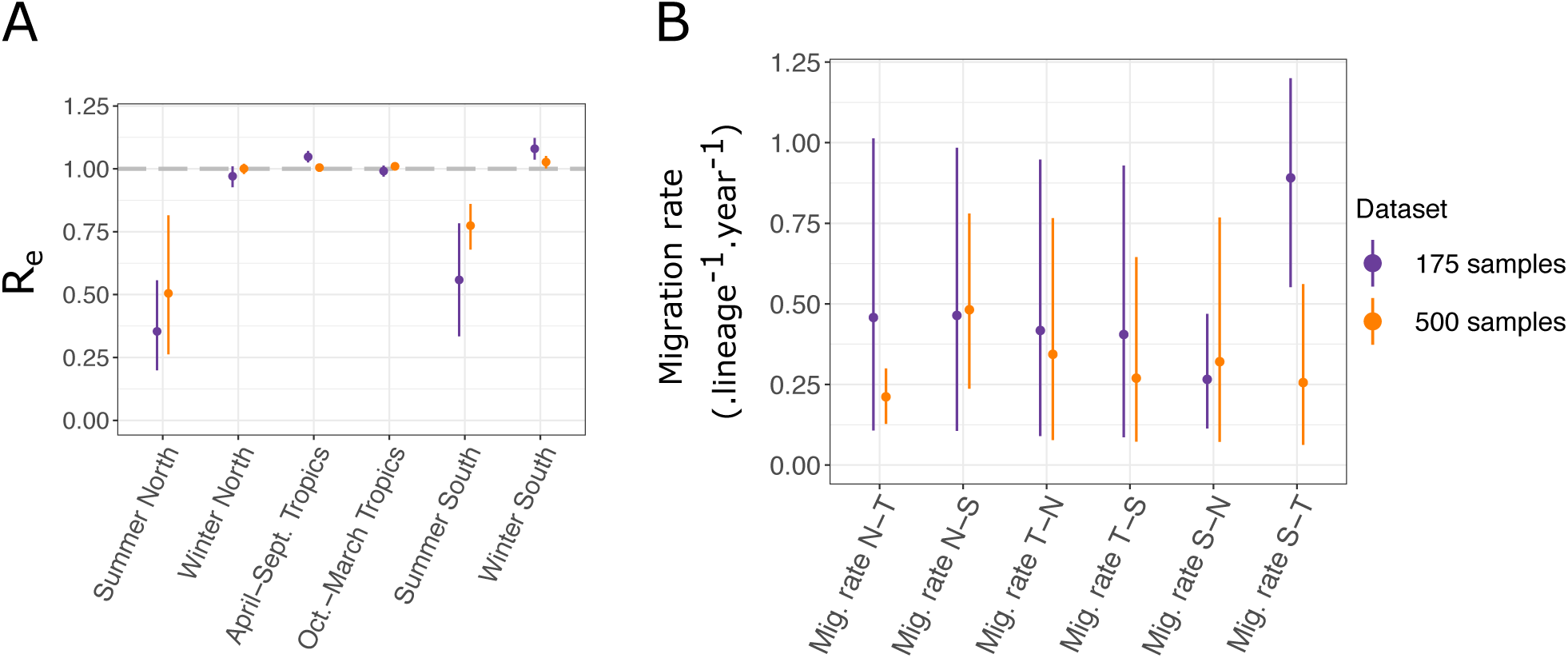
**A:** Seasonal effective reproduction numbers for each sample location, for both datasets. **B:** Migration rates inferred for each dataset. N, T and S refer respectively to North, South and Tropics. For instance, “*Mig. rate N-T*” represents the migration rate from the Northern location to the Tropical one.

Precise inference of migration rates is more difficult, as is reflected by the significant uncertainty we obtain on the estimates (Fig 4B). Still, we observe in general that the uncertainty is reduced for the inference performed with the larger dataset, as expected. A significant difference between the migration rates inferred from the Southern to Tropical locations arises between the two analysis. With the larger dataset, the estimated rate is much lower than with the smaller one, and more in range with the other migration rate estimates. Detailed results of all the parameter estimates for both analyses are available in Table S3. Most notably, estimates of root locations for both datasets are very similar. In both cases, the tropical location is the most likely location for the root; however, neither of the two other locations can be entirely excluded.

## Discussion

The multi-type birth-death model with its updated implementation in the *bdmm* package for BEAST 2 provides a flexible method for taking into account the effect of population structure when performing phylodynamic genetic sequence analysis. Compared to the original implementation, we now prevent underflow of numerical calculations and speed up calculations by almost an order of magnitude. The size limit of around 250 samples for datasets which could be handled by *bdmm* is thus lifted and significantly larger datasets can now be analysed. We demonstrate this improvement by analysing two datasets of Influenza A virus H3N2 genetic data from around the globe. One dataset has 500 samples and could not have been analyzed with the original version of *bdmm*, the other one contains 175 samples and is the original example dataset analyzed in [Kühnert et al., 2016]. Overall, we observe, as could be expected, that analysing a dataset with more samples gives a more long-term picture of the global transmission patterns and reduces the general uncertainty on parameter estimates.

With the addition of so-called *ρ*-sampling events in the past, intense sampling efforts limited to short periods of time (leading to many samples being taken nearly simultaneously) can be easily modelled as instantaneous sampling events across the entire population (or sub-population), rather than as non-instantaneous sampling over small sampling intervals. This simplifies the modelling of intense pathogen sequencing efforts in very short time windows. When using a multi-type birth-death model in the macroevolutionary framework, *ρ*-sampling can be used to model fossil samples originating from the same rock layer. By allowing the removal probability r (the probability for an individual to be removed from the infectious population upon sampling) to be type-dependent and to vary across time intervals, as well as allowing migration rates between types to vary across time intervals, we further increase the generality and flexibility of the multi-type birth-death model.

We focused on an epidemiological application of *bdmm*, where we co-infer the phylogenetic trees to take into account the phylogenetic uncertainty. However, the *bdmm* modelling assumptions are equally applicable to the analysis of macroevolutionary data, in which context *bdmm* allows for the joint inference of trees with fossil samples under structured models. In the context of the exploration of trait-dependent speciation, structured birth-death models such as the binary-state speciation and extinction model (BiSSE) [Maddison, 2006, FitzJohn, 2012] have been shown to possibly produce spurious associations between character state and speciation rate when applied to empirical phylogenies [Rabosky and Goldberg, 2015]. When used in this fashion, users of *bdmm* should assess the propensity for Type I errors with their dataset through neutral trait simulations, as suggested by Rabosky and Goldberg [2015].

In summary, we expect the new release of *bdmm* to become a standard tool for phylodynamic analysis of sequencing data and phylogenetic trees from structured populations.

## Acknowledgements

We thank Nicola Müller, David Rasmussen, Jūlija Pečerska for very valuable discussions and Fábio Kuriki Mendes for helpful comments on the manuscript.

