## Supplementary Information for "Improved multi-type birth-death phylodynamic inference in BEAST 2"

### S1: Derivation of the probability density of a sampled tree

#### S1.1 Probability of having no sampled descendants

In order to compute the probability density of a sampled tree, we need to calculate the probability of an individual with type $i \in \{1 \dots d\}$ at time $t_{k-1} \leq t \leq t_k$ to have no sampled descendant, which we denote by $p_{i,k}(t)$. In the interval $t_{k-1} \leq t < t_k$, this probability satisfies the ordinary differential equation (ODE)

$$\begin{aligned} -\frac{d}{dt}p_{i,k}(t) = &- \left(\sum_{j=1}^{d}(\lambda_{ij,k} + m_{ij,k}) + \mu_{i,k} + \psi_{i,k}\right) p_{i,k}(t) \\ &+ \sum_{j=1}^{d} \lambda_{ij,k} p_{i,k}(t) p_{j,k}(t) + \sum_{j=1}^{d} m_{ij,k} p_{j,k}(t) + \mu_{i,k}. \end{aligned} \tag{S1.1}$$

The terms on the right-hand side in the first line correspond to the probability of no event happening and the terms in the second line to either a branching, migration or death event happening. For a detailed derivation via Master equations, refer to Maddison et al. [2007] or Stadler and Bonhoeffer [2013].

For $t = t_k$, $k \in \{0 \dots n\}$, we have,

$$p_{i,k}(t_k) = (1 - \rho_{i,k}) \times \begin{cases} 1 & \text{if } k = n, \\ p_{i,k+1}(t_k) & \text{otherwise.} \end{cases} \tag{S1.2}$$

#### S1.2 Probability density of a sample-typed subtree

We use $g_{i,k}^{e}(t)$ to denote the probability density that an individual represented by branch $e$ at time $t$ (with $t_{k-1} \leq t \leq t_k$, $k \in \{1 \dots n\}$) in state $i \in \{1, \dots, d\}$ has evolved between $t$ and $T$ as observed in the tree. Branch $e$ is connected to two nodes, and we denote the more recent node with $n_e$, occurring at time $t_{k-1} < t_e \leq t_k$. Node $n_e$ represents either a sampling event (Fig. S1A), a branching event (Fig. S1B), or a degree-2 node at $t_e = t_k$ with or without sampling (Fig. S1C and D). One can show that $g_{i,k}^{e}(t)$ for $t < t_e$ satisfies the ODE

$$
\begin{aligned}
-\frac{d}{dt}g^{e}_{i,k}(t) = &-\left(\sum_{j=1}^{d}(\lambda_{ij,k}+m_{ij,k})+\mu_{i,k}+\psi_{i,k}\right)g^{e}_{i,k}(t) \\
&+\sum_{j=1}^{d}m_{ij,k}g^{e}_{j,k}(t)+\sum_{j=1}^{d}\lambda_{ij,k}p_{j,k}(t)g^{e}_{i,k}(t)+\sum_{j=1}^{d}\lambda_{ij,k}p_{i,k}(t)g^{e}_{j,k}(t),
\end{aligned}
\tag{S1.3}
$$

with the derivation being analogous to that of $p_{i,k}(t)$.

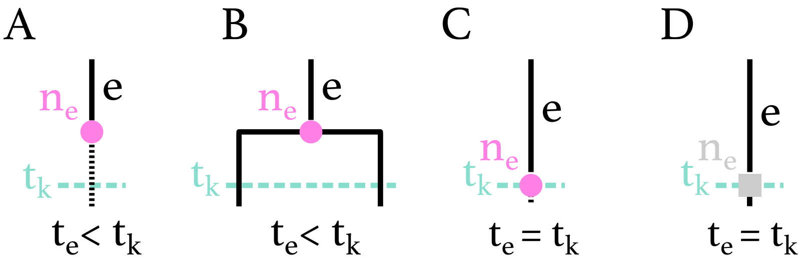


**Figure S1.** Possible configurations for node $n_e$ on branch $e$ at time $t_{k-1} < t_e \leq t_k$. **A:** $n_e$ is a $\psi$-sampled node at time $t_e < t_k$, with or without sampled descendants. **B:** $n_e$ is a branching event at time $t_e < t_k$. **C:** $n_e$ is a $\rho$-sampling node at time $t_e = t_k$, with or without sampled descendants. **D:** $n_e$ is a degree-2 node at time $t_e = t_k$ without sampling.

We denote the branch (resp. two branches) descending from $n_e$ at time $t_e$ with $e_1$ (resp. $e_1$ and $e_2$). The initial conditions for the differential equations at node $n_e$, i.e. the values of the probability densities at the most recent end of the branch $e$, are as follows:

$$
g^{e}_{i,k}(t_e) = \begin{cases}
\psi_{i,k}(r_{i,k}+(1-r_{i,k})p_{i,k}(t_e)) & \text{if } n_e \text{ is a } \psi\text{-sampled tip of type } i, \\
\psi_{i,k}(1-r_{i,k})g^{e_1}_{i,k}(t_e) & \text{if } n_e \text{ is a } \psi\text{-sampled ancestor of type } i, \\
\rho_{i,k}(r_{i,k}+(1-r_{i,k})p_{i,k+1}(t_e)) & \text{if } n_e \text{ is a } \rho\text{-sampled tip of type } i, \\
\rho_{i,k}(1-r_{i,k})g^{e_1}_{i,k+1}(t_e) & \text{if } n_e \text{ is a } \rho\text{-sampled ancestor of type } i, \\
0 & \text{if } n_e \text{ is a sample of type } j \neq i, \\
(1-\rho_{i,k})g^{e_1}_{i,k+1}(t_e) & \text{if } n_e \text{ is not a sample and } t_e = t_k, \\
\frac{1}{2}\sum_{j=1}^{d}\lambda_{ij,k}\left[g^{e_1}_{i,k}(t_e)g^{e_2}_{j,k}(t_e)+g^{e_1}_{j,k}(t_e)g^{e_2}_{i,k}(t_e)\right] & \text{if } n_e \text{ has two descendant branches.}
\end{cases}
\tag{S1.4}
$$

The $\frac{1}{2}$ in the last equation is needed since we compute the probability density of an oriented tree.

#### S1.3 Probability density of a branch-typed subtree

When inferring branch-typed trees, we condition on the type of a branch at all times. Thus, we do not integrate over migration events or unobserved branching events changing the type of the tree lineage. We define $\eta_{i,j} = 1$ for $i \neq j$ and $\eta_{i,j} = 2$ for $i = j$. Equation S1.3 is replaced by,

$$-\frac{d}{dt} g^e_{i,k}(t) = -\left(\sum_{j=1}^{d} (\lambda_{ij,k} + m_{ij,k}) + \mu_{i,k} + \psi_{i,k}\right) g^e_{i,k}(t) + \sum_{j=1}^{d} \eta_{i,j} \lambda_{ij,k} p_{j,k}(t) g^e_{i,k}(t), \quad \text{(S1.5)}$$

with initial conditions

$$g^e_{i,k}(t_e) = \begin{cases} \psi_{i,k}\left(r_{i,k} + (1 - r_{i,k})\, p_{i,k}(t_e)\right) & \text{if } n_e \text{ is a } \psi\text{-sampled tip of type } i, \\ \psi_{i,k}\,(1 - r_{i,k})\, g^{e_1}_{i,k}(t_e) & \text{if } n_e \text{ is a } \psi\text{-sampled ancestor of type } i, \\ \rho_{i,k}\left(r_{i,k} + (1 - r_{i,k})\, p_{i,k+1}(t_e)\right) & \text{if } n_e \text{ is a } \rho\text{-sampled tip of type } i, \\ \rho_{i,k}\,(1 - r_{i,k})\, g^{e_1}_{i,k+1}(t_e) & \text{if } n_e \text{ is a } \rho\text{-sampled ancestor of type } i, \\ 0 & \text{if } n_e \text{ is a sampled tip of type } j \neq i, \\ (1 - \rho_{i,k})\, g^{e_1}_{i,k+1}(t_e) & \text{if } n_e \text{ is not a sample and } t_e = t_k, \\ m_{ij,k}\, g^{e_1}_{j,k}(t_e) + \lambda_{ij,k}\, g^{e_1}_{j,k}(t_e)\, p_{i,k}(t_e) & \text{if } n_e \text{ has one descendant branch with type } j \neq i, \\ \frac{1}{2}\eta_{i,j}\lambda_{ij,k}\, g^{e_1}_{i,k}(t_e)\, g^{e_2}_{j,k}(t_e) & \text{if } n_e \text{ has two descendant branches.} \end{cases} \quad \text{(S1.6)}$$

Here also, the $\frac{1}{2}$ in the last equation is needed since we compute the probability density of an oriented tree. In branch-typed trees, a branch is always of one single type. Given the type of branch $e$ is $i$ implies that $g^e_{j,k}(t) = 0$ for $j \neq i$. Indeed, Eqn. S1.6 states $g^e_{j,k}(t_e) = 0$ for $i \neq j$. Further, Eqn. S1.5 specifies $g^e_{j,k}(t) = 0$ for all $t < t_e$. Note that our current implementation of the branch-typed tree model inference algorithm does not allow birth events giving rise to different types, meaning we assume $\lambda_{i,j} = 0$ for $i \neq j$ when branch-typed trees are used.

#### S1.4 Probability density of a sampled tree

The probability density of a sampled tree, with the lineage at time $t = 0$ being of type $i$ and the branch being labelled with $e$, is the product of the probability density that the individual evolved as observed in the tree ($g^e_{i,1}(0)$ and the probability $h_i$ that the individual at the start of the process is in type $i$.

Hence, the probability density of an oriented sampled tree $\mathcal{T}$ under the multi-type birth–death model is

$$f(\mathcal{T}|\boldsymbol{\lambda}, \boldsymbol{\mu}, \boldsymbol{\psi}, \boldsymbol{m}, T) = \sum_{i=1}^{d} h_i \; g^e_{i,1}(0).$$

In BEAST 2, we infer labelled sampled trees, thus we need to calculate the probability of a labelled sampled tree. In a labelled sampled tree, each sample has a unique label, and orientations at branching events are ignored. In order to obtain the probability density of a labelled sampled tree, we need to transform the oriented tree probability density by multiplying by $2^M/N!$ where $M$ is the number of branching events in the tree and $N$ is the number of samples [Gavryushkina et al., 2014]. The probability density of a labelled tree $\mathcal{T}$ under the multi-type birth–death model is thus

$$\tilde{f}(\mathcal{T}|\boldsymbol{\lambda}, \boldsymbol{\mu}, \boldsymbol{\psi}, \boldsymbol{m}, T) = \frac{2^M}{N!} \sum_{i=1}^{d} h_i \; g^e_{i,0}(0). \qquad \text{(S1.7)}$$

### S2: Improved implementation of the tree probability density evaluation

The core component of the *bdmm* package is the evaluation of the tree probability density given by Eq. (S1.7). This involves numerical integration to solve the system of ODEs defined by Eqs. (S1.1)–(S1.2) and either Eqs. (S1.3)–(S1.4) (for sample-typed trees) or Eqs. (S1.5)–(S1.6) (for branch-typed trees).

In what follows, we discuss the numerical stability of our calculations.

#### S2.1 Extended numerical representation

The traditional floating point representation of a real number $x$ is the closest number $\hat{x}$ to $x$, where

$$\hat{x} = m \times 2^q \tag{S2.8}$$

and where $m$ (the *mantissa*) and $q$ (the *exponent*) are signed binary integers of a specified number of bits. Note that the mantissa is understood to represent a fixed-precision number between 1 and 2. In the standard 64 bit double-precision floating point (DPFP) used by the original *bdmm* implementation [Kühnert et al., 2016], $m$ is restricted to 53 bits and $q$ is restricted to 11 bits. Ignoring the mantissa, this implies that the smallest representable non-zero absolute value is on the order of $2^{-2^{10}} \simeq 10^{-320}$. We calculate a probability density $\tilde{f}$ over tree space (together with intermediate values $g^e_{i,k}(t)$ as part of the ODE integration), and these values can fall well below the lower limit imposed by the standard 64 bit DPFP representation, resulting in underflow errors.

To work around this limitation, our new implementation of *bdmm* employs a 96 bit extended precision floating point representation (EPFP) in which the mantissa is represented using a standard 64 bit double precision floating point number and the exponent is represented using a 32 bit signed integer value. This dramatically extends the range of possible values, in particular the smallest non-zero absolute value is $2^{-2^{31}} \simeq 10^{-646456993}$.

A naive approach to employing the EPFP representation would be to use numbers of this kind at each and every step in the probability density calculation. This would ensure that all intermediate calculation results were accurately stored and would allow for accurate calculations at the next step.

However, this approach has two main drawbacks. Firstly, existing numerical integration libraries almost exclusively use the DFPF representation. While integration algorithms could certainly be implemented for other representations such as the EPFP, this would be extremely time-consuming. Secondly, since DPFP calculations are implemented at a hardware level on modern processors, calculations using this representation are done very efficiently. Abandoning this primitive data type thus makes basic calculation steps considerably less efficient.

For these reasons, our approach involves a mixture using both representations. Numerical integration of the ODEs along each branch of the sampled phylogeny (Eqns. S1.3 and S1.5) is performed using the DPFP representation, while the combination of these integration results – done to provide the boundary condition for the next integration (Eqns. S1.4 and S1.6) – is achieved using the EPFP representation.

In order to avoid underflow at all times, the EPFP numbers are scaled up before they are converted to the DPFP representation used by the integrator. The goal here is to make sure all values stored as DPFP actually fit the window of values this representation can express. After each numerical integration step, results are converted back to EPFP and rescaled accordingly.

This whole process amounts to an integration by substitution. We scale $g^e_{i,k}(t_k)$ by the linear substitution function $G^e_{i,k}(t_k) = \delta \times g^e_{i,k}(t_k)$, with $\delta$ being a scale factor. Since the original equations (S1.3) and (S1.5) are both linear in $g^e_i$, this scaling does not affect the results of their integration once the inverse scaling at time $t_{k-1}$ is done. The method employed to choose an appropriate scale factor is described in detail below. The differential equation for the probability $p_{i,k}(t)$ of having no sampled descendants is not scaled at all as its integration does not cause any underflows in practice.

#### S2.2 Choice of a scale factor

The scale factor is shared among all subpopulations. Therefore, it has to be carefully chosen, so that all initial conditions fit in the window of values that can be represented in DPFP. We use three rules, A, B, C, when looking for an appropriate scale factor. We apply rule A if possible, otherwise rule B, otherwise rule C. The rules are illustrated in Figure S2 and stated in words in the corresponding legend.

#### S2.3 Performance improvements

We use various techniques to increase the efficiency of the numerical calculations performed by *bdmm*.

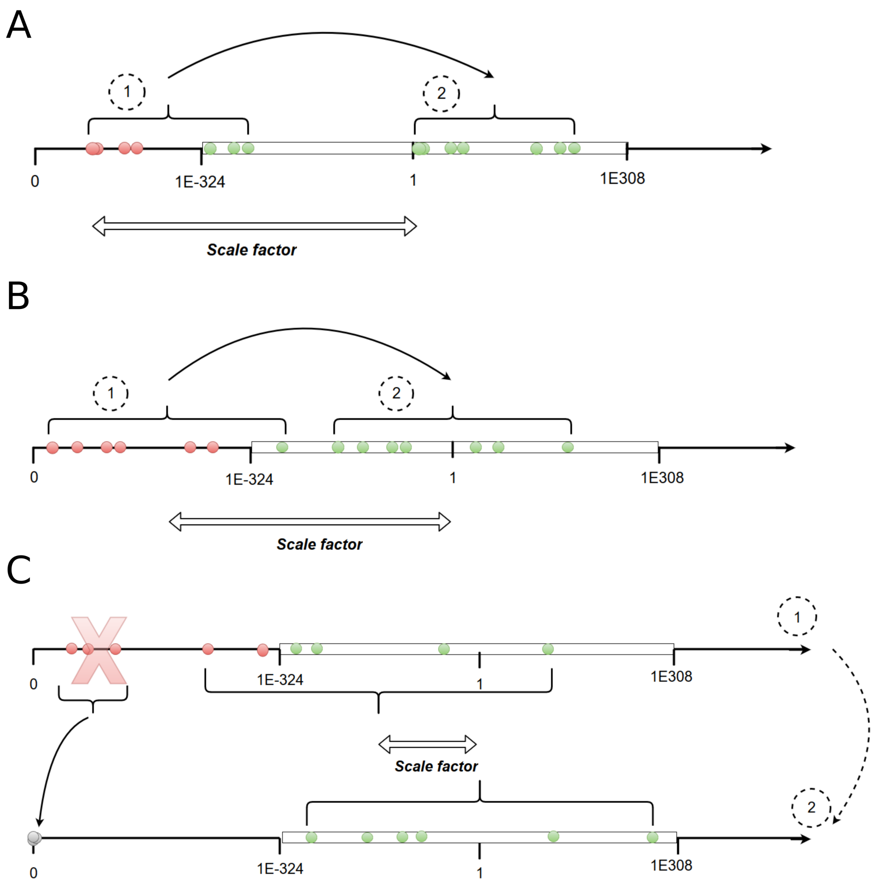


**Figure S2.** Scale factor choice. **A:** Simplest case. The scale factor is the inverse of the smallest non-scaled value. **B:** If A is not applicable, the scale factor is chosen such that the initial conditions are centered inside the range of values acceptable. The mid-point (on a log-scale) of this interval is approximately 1. **C:** Last case, if all scaled values cannot fit at once inside the range of accepted values, the lowest non-scaled values are dropped and set to zero so that the problem simplifies to case 1 (panel A) or 2 (panel B). In all panels, the white rectangle represents values that can be represented using DPFP. Dots represents the values of initial conditions for the differential equations of the multi-type birth-death model, before (1) and after (2) scaling. Red dots represent values that are initially outside the window of values that can be represented using DPFP.

Prior to the integration of the coupled differential equations $p_{i,k}$ and $g^e_{i,k}$ backwards in time, we calculate the values of $p_{i,k}$ for every sampling event in the tree, by numerically integrating the ODEs for $p_{i,k}$. We store the values obtained and use them when calculating the initial conditions for $g^e_{i,k}$ for every tip.

We also implement a parallelization scheme. An initial recursive tree traversal step is necessary to reach the tips of the tree, before launching numerical integration of the system of ODEs on $g^e_{i,k}$ and $p_{i,k}$ along the tree branches. During this traversal, when a node is reached whose left and right child subtrees are of significant size compared to the total tree size, a new computation thread is spawned and assigned to the traversal of one of the two subtrees. The initial thread continues onward with the traversal of the other one. This split between two threads is only executed when both subtrees represent more than a user-defined fraction of the total tree length, by default a tenth. This is done to prevent excessive numbers of threads from being created, since thread creation itself carries a computational overhead.

Finally, we replace the fixed-step size Runge-Kutta integrator used as default integrator in the original implementation by the fifth-order Dormand-Prince integrator for ODEs [Dormand and Prince, 1980]. This integrator uses stepsize control, which improves the efficiency and accuracy of numerical integration steps. We use the existing implementation of these integrators from the *Apache Commons Math* Java library.

#### S3: Additional details on influenza data analysis

As we deal with pathogen sequence data, we adopt the epidemiological parametrization of the multi-type birth-death model as detailed in Kühnert et al. [2016]. The epidemiological parametrization substitutes birth, death and sampling rates with effective reproduction numbers within types, rate at which hosts become noninfectious and sampling proportions. For $i \in \{1, \dots, d\}$ and $k \in \{1, \dots, n\}$,

$$R_{i,k} = \frac{\lambda_{ii,k}}{\mu_{i,k} + r_{i,k}\,\psi_{i,k}}.$$

The rate of becoming uninfectious $\delta_{i,k}$ represents the inverse of the mean duration of infection:

$$\delta_{i,k} = \mu_{i,k} + r_{i,k}\,\psi_{i,k}.$$

Based on our data, $\rho_{i,k} = 0$ for all $i, k$, as there is no singular point in time when a population-wide sampling effort was carried out which lead to multiple simultaneous samples. Thus, the probability for an individual to be sampled, or sampling proportion, is:

$$s_{i,k} = \frac{\psi_{i,k}}{\mu_{i,k} + \psi_{i,k}}.$$

We assume $r = 1$ for all $i, k$ in our analyses, i.e. individuals become non-infectious upon sampling. Further, we assume that $\delta$ is constant across locations $i$ and time intervals $k$. Sampling proportion and migration are assumed not to change through time.

To study the seasonal dynamics of the global epidemic, we allow the effective reproduction number $R$ to vary through time. To do so, we subdivide time into six-month intervals (starting April 1st and October 1st) for the time period during which we have samples. We set all values corresponding to the same season across different years to be equal for a particular location. Therefore, we infer two different values of $R$ for each location $X$, one which corresponds to the April-September period: $R_{X_1}$ and another one for the October-March period: $R_{X_2}$. The location-specific sampling proportions $s_i$ are assumed to be constant in the time interval in which we have samples, and null before the first sample. Migration rates between types are inferred as products of a unique migration factor $\sigma_m$ and relative rates $M_{i,j}$:

$$\forall\, i, j \in \{1 \dots d\}, m_{i,j} = \sigma_m \, M_{i,j}.$$

This setting allows us to use rather informative priors around 1 for the relative rates, and only requires a less informative prior for the unique migration factor. Supplementary Table 1 lists the prior distributions assumed for the birth-death parameters.

Following Vaughan et al. [2014], we use a GTR+$\Gamma$ substitution model [Lanave et al., 1984] and a strict molecular clock with the same priors as in Vaughan et al. [2014].

The BEAST 2 analysis infers the birth-death model parameters, the substitution model parameters, and the clock model parameters together with the phylogenetic tree. In our analysis, the inferred phylogenetic trees are sample-typed trees. Thus, we do not attempt to reconstruct the history of migrations, rather we marginalize over all the possible migration histories. This is done to allow the MCMC chain to converge in shorter time compared to an analysis inferring branch-typed trees.

For each data analysis, we run 10 parallel MCMC chains for $9 \times 10^7$ steps each using the new implementation of *bdmm* as a package of BEAST 2.6 [Bouckaert et al., 2014, 2019]. We use Tracer v1.6 [Rambaut et al., 2018] to check for convergence. We combine the chains run in parallel into one using LogCombiner. We obtain the MCC tree using

TreeAnnotator. Finally, we plot the numerical results with the R package *ggplot2* [Wickham, 2016] and the MCC tree with *ggtree* [Yu et al., 2017]. The effective sample size was above 200 for all parameters.

### S4: Supplementary tables

| Parameter | Distribution |
|---|---|
| $\lambda_i$ | Unif(1, 3) |
| $\mu_i$ | $\lambda_i \times$ Unif(0, 1) |
| $m_{i,j}$ | Unif(0, 0.5) |
| $\psi_i$ | Unif(0.05, 0.5) |
| $r_i$ | Unif(0, 1) |

**Table S1.** Distributions from which parameters were sampled for simulating trees. All parameters are constant through time.

| Parameter | Prior distribution |
|---|---|
| $R$ | LogNormal(0, 1.0) |
| $\delta$ | LogNormal(4.5, 0.15) |
| $\sigma_m$ | LogNormal(0, 2.0) |
| $M_{i,j}$ | LogNormal(0, 0.5) |
| $s$ | Exp(0.001) truncated on $[0\,,1]$ |
| $r$ | Beta(10.0, 1.5) |
| $T$ | LogNormal(2.0, 1.0) |

**Table S2.** Prior distributions for parameters of the multi-type birth-death model in the seasonal influenza analysis.

| | 175 samples | | | 500 samples | | |
|---|---|---|---|---|---|---|
| | m | hpd_low | hpd_high | m | hpd_low | hpd_high |
| $t$ | 3.342 | 3.048 | 3.643 | 6.645 | 6.313 | 7.031 |
| $\delta$ | 90.464 | 75.836 | 105.207 | 101.582 | 87.197 | 116.768 |
| $R_{N_s}$ | 0.354 | 0.199 | 0.556 | 0.505 | 0.263 | 0.815 |
| $R_{N_w}$ | 0.971 | 0.927 | 1.01 | 1.001 | 0.98 | 1.02 |
| $R_{T_{as}}$ | 1.048 | 1.026 | 1.071 | 1.005 | 0.992 | 1.017 |
| $R_{T_{om}}$ | 0.991 | 0.969 | 1.013 | 1.01 | 0.998 | 1.022 |
| $R_{S_s}$ | 0.558 | 0.335 | 0.783 | 0.774 | 0.679 | 0.861 |
| $R_{S_w}$ | 1.08 | 1.037 | 1.123 | 1.027 | 1.002 | 1.051 |
| $\sigma_m$ | 0.475 | 0.196 | 0.869 | 0.304 | 0.147 | 0.524 |
| $M_{N,T}$ | 0.871 | 0.245 | 1.923 | 1.064 | 0.3 | 2.422 |
| $M_{N,S}$ | 0.894 | 0.253 | 1.965 | 0.838 | 0.264 | 1.744 |
| $M_{T,N}$ | 2.183 | 0.877 | 4.174 | 1.568 | 0.635 | 2.973 |
| $M_{T,S}$ | 0.561 | 0.205 | 1.08 | 0.694 | 0.332 | 1.248 |
| $M_{S,N}$ | 1.001 | 0.273 | 2.261 | 0.887 | 0.275 | 1.93 |
| $M_{S,T}$ | 1.005 | 0.279 | 2.173 | 1.13 | 0.368 | 2.468 |
| $h_N$ | 0.292 | 0 | 0.777 | 0.296 | 0 | 0.779 |
| $h_T$ | 0.3 | 0 | 0.784 | 0.292 | 0 | 0.777 |
| $h_S$ | 0.283 | 0 | 0.769 | 0.289 | 0 | 0.767 |
| $P$(root in N) | 0.024 | 0 | 0.535 | 0.023 | 0 | 0.501 |
| $P$(root in T) | 0.849 | 0.167 | 1 | 0.863 | 0.176 | 1 |
| $P$(root in S) | 0.041 | 0 | 0.731 | 0.044 | 0 | 0.67 |

**Table S3.** Inferred parameter values for Influenza A virus analysis under the multi-type birth-death model. For each parameter, the lower and upper bounds for the 95% Highest Posterior Density interval (hpd_low, hpd_high) are given along with the median (m). $N$, $T$, and $S$ refer respectively to the North, South and Tropics. For effective reproduction numbers $R$, the first subscript is the location while the second one refers to the period of the year. $s$, $w$, $as$, $om$ respectively refer to *summer*, *winter*, *april-september*, and *october-march.* Thus, for instance, $R_{S_w}$ refers to the effective reproduction number of samples from the southern hemisphere during the winter season. The tree height $t$ is given in number of years, $M$ is given in migrations/lineage/year and $\delta$ is given in years$^{-1}$. The remaining parameters are unitless.

Remco Bouckaert, Timothy G. Vaughan, Joëlle Barido-Sottani, Sebastián Duchêne, Mathieu Fourment, Alexandra Gavryushkina, Joseph Heled, Graham Jones, Denise Kühnert, Nicola De Maio, Michael Matschiner, Fábio K. Mendes, Nicola F. Müller, Huw A. Ogilvie, Louis du Plessis, Alex Popinga, Andrew Rambaut, David Rasmussen, Igor Siveroni, Marc A. Suchard, Chieh-Hsi Wu, Dong Xie, Chi Zhang, Tanja Stadler, and Alexei J. Drummond. Beast 2.5: An advanced software platform for bayesian evolutionary analysis. *PLOS Computational Biology*, 15(4):1–28, 04 2019. doi: 10.1371/journal.pcbi.1006650. URL `https://doi.org/10.1371/journal.pcbi.1006650`.

John R Dormand and Peter J Prince. A family of embedded runge-kutta formulae. *Journal of computational and applied mathematics*, 6(1):19–26, 1980.

Alexandra Gavryushkina, David Welch, Tanja Stadler, and Alexei J Drummond. Bayesian inference of sampled ancestor trees for epidemiology and fossil calibration. *PLoS computational biology*, 10(12):e1003919, 2014.

Denise Kühnert, Tanja Stadler, Timothy G. Vaughan, and Alexei J. Drummond. Phylodynamics with Migration: A Computational Framework to Quantify Population Structure from Genomic Data. *Molecular Biology and Evolution*, 33(8):2102–2116, 2016. ISSN 15371719. doi: 10.1093/molbev/msw064.

Cecilia Lanave, Giuliano Preparata, Cecilia Sacone, and Gabriella Serio. A new method for calculating evolutionary substitution rates. *Journal of Molecular Evolution*, 20(1): 86–93, 1984. URL `https://link.springer.com/article/10.1007/BF02101990`.

Wayne P Maddison, Peter E Midford, and Sarah P Otto. Estimating a binary character's effect on speciation and extinction. *Systematic biology*, 56(5):701–10, 2007. ISSN 1063-5157. doi: 10.1080/10635150701607033. URL http://www.ncbi.nlm.nih.gov/pubmed/17849325.

Andrew Rambaut, Alexei J Drummond, Dong Xie, Guy Baele, and Marc A Suchard. Posterior Summarization in Bayesian Phylogenetics Using Tracer 1.7. *Systematic Biology*, 67(5):901–904, 04 2018. ISSN 1063-5157. doi: 10.1093/sysbio/syy032. URL https://doi.org/10.1093/sysbio/syy032.

Tanja Stadler and Sebastian Bonhoeffer. Uncovering epidemiological dynamics in heterogeneous host populations using phylogenetic methods. *Philosophical transactions of the Royal Society of London. Series B, Biological sciences*, 368(1614):20120198, 2013. ISSN 1471-2970. doi: 10.1098/rstb.2012.0198. URL http://www.ncbi.nlm.nih.gov/pubmed/23382421.

Timothy G. Vaughan, Denise Kühnert, Alex Popinga, David Welch, and Alexei J. Drummond. Efficient Bayesian inference under the structured coalescent. *Bioinformatics*, 30(16):2272–2279, 04 2014. ISSN 1367-4803. doi: 10.1093/bioinformatics/btu201. URL https://doi.org/10.1093/bioinformatics/btu201.

Hadley Wickham. *ggplot2: Elegant Graphics for Data Analysis*. Springer-Verlag New York, 2016. ISBN 978-3-319-24277-4. URL https://ggplot2.tidyverse.org.

Guangchuang Yu, David K. Smith, Huachen Zhu, Yi Guan, and Tommy Tsan-Yuk Lam. ggtree: an r package for visualization and annotation of phylogenetic trees with their covariates and other associated data. *Methods in Ecology and Evolution*, 8(1):28–36, 2017. doi: 10.1111/2041-210X.12628. URL https://besjournals.onlinelibrary.wiley.com/doi/abs/10.1111/2041-210X.12628.
